# Combining interstrand crosslinking agents with histone deacetylase inhibitors against high grade IDH mutant gliomas

**DOI:** 10.64898/2026.09.28.755068

**Authors:** Thomas K. Sears, Lillian Coe, Rahul Chaliparambil, Wenxia Wang, Mateo Gomez, Matthew McCord, Andrew Li, Susan Gueble, Ranjit Bindra, Craig Horbinski

## Abstract

Mismatch repair (MMR) deficiency contributes to temozolomide (TMZ) resistance in a subset of *isocitrate dehydrogenase* 1/2 mutant (IDH^mut^) gliomas, creating a need for therapies that retain activity despite MMR loss. We evaluated the novel interstrand crosslinking agent KL-50, alone and in combination with the histone deacetylase inhibitor belinostat, in patient-derived IDH^mut^ glioma models. MSH6 knockout or constitutive expression established four matched MMR-proficient and MMR-deficient culture models. Drug responses were assessed using live cell count and cytotoxicity assays, and KL-50 monotherapy was tested in intracranial BT142 orthotopic xenografts. LOEWE synergy analysis, RNA sequencing, and a genome-wide CRISPR knockout screen assessed combination activity and potential mechanisms. MMR deficiency increased TMZ GI_₅₀_ values 2.2- to 30.4-fold, whereas KL-50 retained antiproliferative activity and showed enhanced cytotoxicity in selected MMR-deficient models. In TMZ-naïve, MMR-proficient BT142 xenografts, KL-50 increased median survival from 107.5 to 194 days. Two of eight treated mice had no histologically detectable residual tumor after 220 days post-engraftment. KL-50 and belinostat exhibited synergistic cytotoxicity in BT142, 905, and TS603 IDH^mut^ glioma cultures. Transcriptomic gene ontology analyses identified downregulation of DNA repair pathways following histone deacetylase inhibition, including terms for homologous recombination and double-strand break repair. The CRISPR screen identified depletion of PKMYT1-targeting gRNAs during KL-50 treatment, implicating this mitotic checkpoint kinase as a candidate determinant of drug sensitivity. PKMYT1 expression was also reduced by histone deacetylase inhibition. These findings support further preclinical development of KL-50, alone and in combination with belinostat, for IDH^mut^ gliomas including tumors with MMR-associated TMZ resistance.

## Introduction

Mutations in *isocitrate dehydrogenase 1* or *2* (IDH^mut^) define two major adult-type diffuse glioma lineages: astrocytoma, IDH^mut^, CNS WHO grades 2-4, and oligodendroglioma, IDH^mut^ and 1p/19q-codeleted, CNS WHO grades 2-3.(1) IDH^mut^ gliomas are less aggressive than IDH wildtype (IDH^wt^) glioblastoma (GBM), yet they are still diffusely infiltrative, inevitably recurrent, and ultimately fatal. Although the IDH^mut^ enzyme inhibitor vorasidenib is effective against grade 2 minimally enhancing IDH^mut^ astrocytomas and oligodendrogliomas,(2) such inhibitors are much less effective against higher grade IDH^mut^ gliomas.(3,4) Since most IDH^mut^ gliomas either present as higher grade and/or will eventually progress to higher grade even with inhibitor therapy, additional treatment strategies are still needed for this subgroup of brain cancers.

Current standard-of-care for higher grade IDH^mut^ gliomas includes adjuvant treatment with the alkylating agent temozolomide (TMZ). TMZ produces cytotoxic O^6^-methylguanine lesions that are removed by O^6^-methylguanine-DNA methyltransferase (MGMT).(5) Since most IDH^mut^ gliomas silence MGMT expression via promoter methylation,(6) they are more sensitive to TMZ. However, TMZ cytotoxicity still depends on an intact DNA mismatch-repair (MMR) pathway to recognize alkylated DNA and attempt to repair the damage, triggering a futile recognition-repair cycle that eventually leads to apoptosis.(7) Prolonged TMZ exposure often selects for MMR-deficient glioma clones that tolerate O^6^-methylguanine while accumulating extensive TMZ-signature mutations, especially in IDH^mut^ astrocytomas and oligodendrogliomas.(8,9)

KL-50 is a new brain-penetrant imidazotetrazine designed to remain active in both MMR-proficient and -deficient conditions. KL-50 initially produces an MGMT-reversible DNA lesion that, when unrepaired in MGMT-deficient tumor cells, develops into an interstrand cross-link and induces MMR-independent cell death.(10) In IDH^wt^ GBM models, KL-50 is active against both MMR-proficient and MMR-deficient tumors while remaining well tolerated in mice.(10) Subsequent studies by our group in patient-derived IDH^wt^ GBM models confirmed activity against both treatment-naive and recurrent, post-TMZ MMR-deficient gliomas.(11) Despite these promising results with KL-50, monotherapies rarely produce durable cancer control. Combination therapies, based on rational mechanisms of therapeutic additivity or synergy, are more likely to succeed in clinical trials. We and others previously showed that IDH^mut^ increases glioma cell sensitivity to histone deacetylase inhibitors (HDACi), including the FDA-approved HDACi belinostat, resulting in more histone acetylation, growth inhibition, and apoptosis compared to IDH^wt^ GBM cells.(12,13) As an extension of that work, we sought to evaluate KL-50 with and without belinostat against high-grade IDH^mut^ gliomas, including explorations into potential mechanisms of synergy.

## Materials and Methods

### Patient-derived Glioma 3D Spheroid Models

IDH^mut^ patient-derived BT142, 905, MGG119, and MGG152 glioma cultures were acquired from various sources. BT142 was purchased from ATCC, 905 was a gift from Dr. Kevin Woolard at UC Davis, with MGG119 and MGG152 received from Dr. Harley Kornblum at UCLA. All cultures were grown in defined media lacking FBS as previously described.(14) For both BT142 and 905, supplemental PDGF-AA was added to the growth media at a concentration of 100 ng/mL.

### Generation of MSH6 Knockout (KO)/Mismatch Repair-deficient IDH^mut^ Glioma Models

Lentiviral constructs to establish MMR+ and MMR-glioma models were purchased from VectorBuilder. For BT142, MGG119, and MGG152, which are MMR-proficient, lentiviral *MSH6* KO was performed using CRISPR methodology. Two *MSH6*-targeting gRNAs (pLV[CRISPR]-hCas9:T2A:Bsd-U6>hMSH6[gRNA#2152]; pLV[CRISPR]-hCas9:T2A:Bsd-U6>hMSH6[gRNA#2158]) were used to induce optimal loss of MSH6 protein levels, and a corresponding gRNA scramble (pLV[CRISPR]-hCas9:T2A:Bsd-U6>Scramble) was used as a control. For 905, which is MMR-deficient, *MSH6* overexpression was performed by using lentivirus-based *MSH6* constitutive expression (pLV[Exp]-Bsd-CMV>hMSH6[NM_001406795.1]) or control lentivirus(pLV[Exp]-EGFP/Puro-EF1A>mCherry), followed by positive antibiotic selection. Validation of *MSH6* knockout or overexpression was assessed using western blots.

### Cell Growth and Cytotoxicity Analysis

A Countess II Live Cell Counter utilizing a trypan blue live/dead stain was employed to evaluate drug-induced changes in cell growth and cytotoxicity. TMZ (SelleckChem), KL-50 (ModifiBio), and belinostat (SelleckChem) were dissolved at 200 mM stock concentrations in DMSO, diluted to 1000x of the target concentrations, and treated with drug or DMSO vehicle at the mentioned doses. Live cell counts and percent live cells were plotted using GraphPad Prism 10.4.2. Doses at which growth was reduced by half (GI50s) were calculated by applying nonlinear regression models to live cell counts.

### Patient-derived Intracranial Xenograft Model of IDH^mut^ glioma

Patient-derived intracranial xenografts were established in NOD *<u>scid</u>*gamma (NSG) mice using TMZ-naïve, MMR-proficient BT142 glioma spheroid cultures. Briefly, BT142 spheroids were trypsinized into a single-cell suspension, counted on our Countess II Live Cell Counter, and then resuspended at a concentration of 25,000 cells/µL in DPBS. Each mouse was then engrafted with 50,000 cells, or 2 µL of cell suspension. All animal procedures aligned with IACUC policies at our institution and were approved by an institutional IACUC review board. Two weeks post-engraftment, mice were initiated on KL-50 at a dose of 10 mg/kg for 3 weeks, daily M-F, via IP administration.

### Intracranial Xenograft H&E Staining and Cell Density Heatmap

Upon reaching a survival endpoint of 15% bodyweight reduction and/or moribund behavior, mice were sacrificed and brain tissue was harvested for histological evaluation via H&E staining under the guidance of a board-certified neuropathologist (CH). Brain tissues were fixed for 24 hr in formalin, sliced into coronal sections, and then placed into cassettes for tissue clearing. Tissue clearing was performed by Northwestern’s Tissue Processing Core. Tissue slices were then paraffin-embedded, sectioned on a microtome, and stained with hematoxylin and eosin by Northwestern’s Nervous System Tumor Biobank (NSTB).

Whole-slide H&E images were acquired at 40× magnification using an Aperio Leica GT450 scanner (0.263 µm/pixel) and analyzed using Python, OpenSlide, and scikit-image.(15,16) Brain tissue was identified at 4.21 µm/pixel using optical-density and color-based thresholding, followed by morphological cleanup. Nuclear quantification was performed in non-overlapping 512 × 512-pixel tiles at 1.05 µm/pixel. Hematoxylin signal was isolated by H&E color deconvolution, with slide-specific intensity thresholding used to account for staining variability. Nuclear density was calculated as the number of detected nuclei per mm² of viable tissue.

A nuclear-density threshold of 3,096 nuclei/mm² was established in a representative pilot slide, confirmed by visual review, and subsequently held constant across all specimens. Areas exceeding this threshold were classified as provisional tumor-like high-cellularity regions. The normally hypercellular cerebellar granule layers in the rightmost posterior section were retained in total tissue-area and nuclear-density measurements but excluded from the tumor-like numerator. The tumor-like area fraction for each mouse was calculated as the area of classified tumor-like tissue divided by the total analyzed brain-tissue area across all sections. Accuracy and regional classification was assessed by comparing whole-slide density maps to representative high-resolution fields.

### Bulk RNA-Seq of Panobinostat- and Belinostat-treated IDH^mut^ Glioma Cultures

IDH^mut^ glioma cultures were harvested after 24 hours of drug treatment (10 nM panobinostat, 500 nM belinostat) and flash-frozen with three biological replicates per condition. RNEasy kits from Qiagen were used for RNA extractions, and a Nanodrop spectrophotometer was used to quantify RNA content. Samples were then submitted to Northwestern’s NUSeq Core for library preparation, sequencing, and data processing. Concerning programs/packages used for analysis, FastQC was used for assessing raw read quality, adapters removed using TrimGalore and CutAdapt, and hg38 genomic alignment applied using STAR.(17,18) Normalized gene counts were calculated using HTSeq and used for downstream differential gene expression (DESeq2) and gene ontology (clusterProfiler) analysis.(19–21)

### CRISPR KO Screen

VectorBuilder’s human whole-genome dual-gRNA library (Cat# LV5M(Lib190505-1046fgb)) was used to perform a CRISPR KO viability screen with KL-50. First, stable Cas9 expression was established in our 905 IDH^mut^ using a Cas9 lentivirus (pLV[Exp]-Bsd-EF1A>hCas9) followed by blasticidin selection. Then, a second transduction was performed at a previously-validated MOI of 0.3, with our whole-genome dual-gRNA pooled lentivirus library targeting 20,048 genes, with 4-6 gRNA sequences per gene. This was then followed by puromycin selection and expansion of cultures for 14 days, with a final KL-50 viability screen conducted for 5 days at a dose of 50 µM. Cells were then counted and harvested, and a minimum final coverage of 125x was calculated for each sample. Due to the large number of cells (>100 million) needed per sample at initial transduction, only two biological replicated were generated per condition. DNA was extracted using a DNEasy Kit (Qiagen), quantified using a Qubit Fluorometer, and sent to VectorBuilder for NGS library preparation, Illumina sequencing, and MAGECK analysis.(22)

### LOEWE Synergy Analysis

BT142, 905, and TS603 IDH^mut^ glioma cultures were treated with KL-50 (10-50 µM), belinostat (100-500 nM), or both drugs for 5 days. Upon treatment completion, cytotoxicity was assessed on a Countess II Live Cell Counter. Cytotoxicity data was prepared in matrix format and input into Combenefit 2.021.(23) For assessment of synergy, a Synergy Score>10 indicates a synergistic effect, which is also distinguished by blue in the heatmap plots.

### Western Blotting

Western blots were performed using Invitrogen’s Novex system with bis-tris gels and MOPS running buffer. Briefly, frozen cell pellets were lysed in RIPA buffer containing 1x EDTA-free Protease Inhibitor Cocktail (Thermo Fisher Scientific, Cat# 89901 and 87785) and then sonicated on a QSonica Q800R3 sonicator. Samples were then cleared of insoluble material via high-speed centrifugation (16,000g) and protein concentrations assessed using Pierce’s Detergent Compatible Bradford Assay (Cat# 23246). Proteins were separated on 4-12% gradient SDS-PAGE gels for 1 hour at 200V, and then and transferred to 0.22 μm PVDF membranes at 30 V for 1 hour. Membranes were blocked with StartingBlock T20 Blocking Buffer (Thermo Fisher Scientific, Cat# 37543) and incubated with primary antibodies diluted 1:1,000 in 5% BSA dissolved in TBST at 4°C overnight, followed by room temperature incubation with respective secondary antibodies (goat anti-rabbit, Cell Signaling Technology #7074; goat anti-mouse, Cell Signaling Technology #7076) in blocking buffer at concentration of 1:10,000 for 60 minutes. Membranes were imaged with SuperSignal West Pico PLUS Chemiluminescent reagents (Thermo Fisher Scientific Cat# 24580) using a BioRad ChemiDoc imaging system. The primary antibodies used in this study were anti-GAPDH (Cat# 2118) and anti-MSH6 (Cat# 5424). An HRP-linked goat anti-rabbit secondary antibody was used for chemiluminescence.

## Results

### Generation of MMR-deficient and -proficient patient-derived IDH^mut^ glioma models via modulation of MSH6 levels

We first evaluated MSH6 protein levels via western blot in a panel of patient-derived IDH^mut^ glioma spheroid cultures. Among IDH^mut^ cells, BT142, MGG119, and MGG152 have high MSH6, whereas 905 lacks MSH6 (**Fig. 1A**). We therefore generated MMR-deficient IDH^mut^ glioma models by lentiviral CRISPR KO of *MSH6* in BT142, MGG119, and MGG152 cells. Two different gRNAs were tested, and MSH6 suppression was achieved with at least one gRNA for each cell source (**Fig. 1B**). Cell derivatives with the greatest reduction in MSH6 protein expression were selected for subsequent experiments. Conversely, the MSH6-deficient 905 cells were transduced with a lentiviral construct to induce MSH6 expression (**Fig. 1C**).

**Figure 1.**
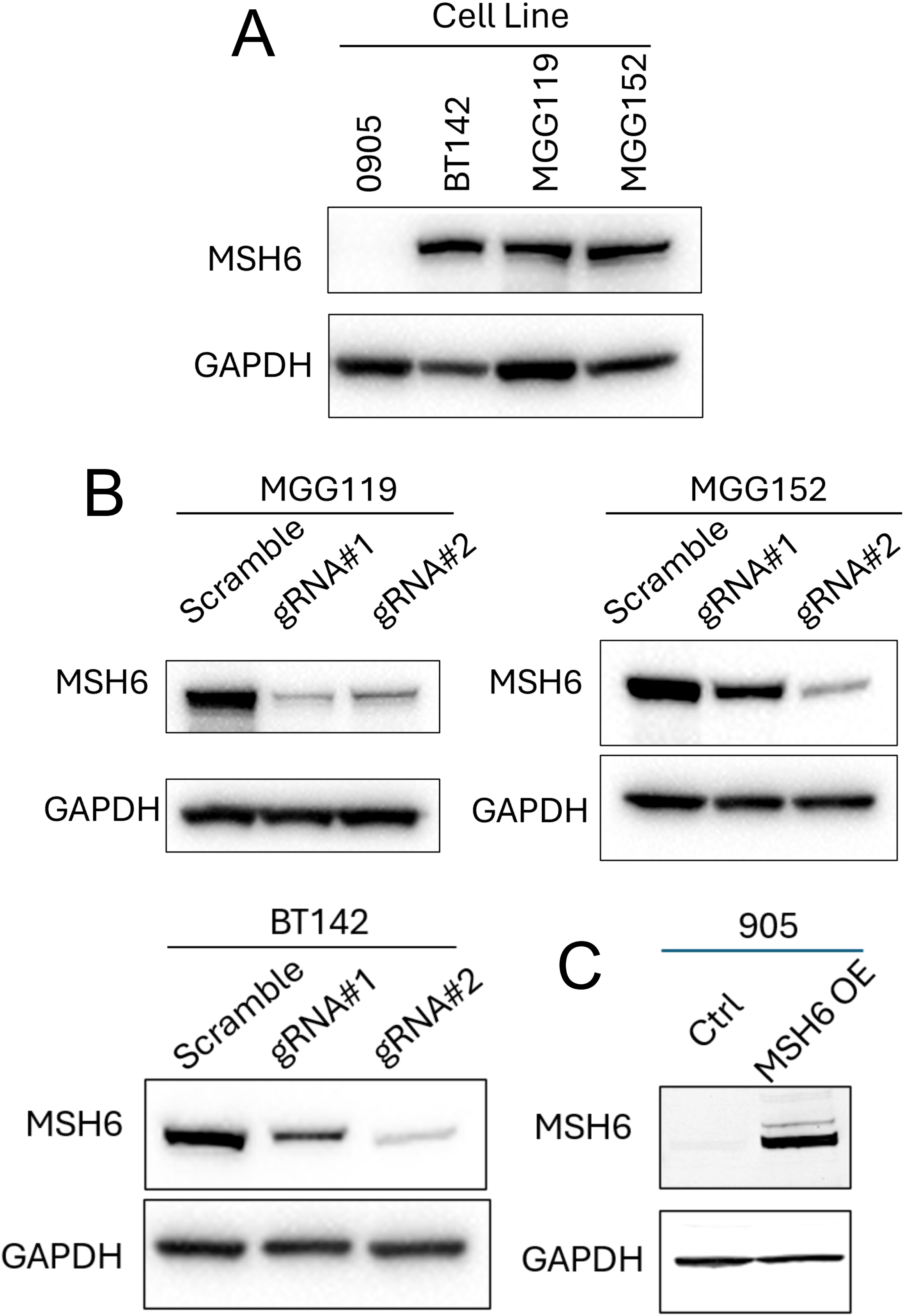
Generation of MMR-proficient and -deficient IDH^mut^ glioma models via MSH6 KO. (A) Baseline expression of MSH6 in our panel of IDH^mut^ glioma cultures. (B) Validation of reduced MSH6 protein levels after CRISPR KO using MGG152, MGG119, and BT142 IDH^mut^ glioma cultures. (C) Validation increased in MSH6 protein levels after constitutive expression of MSH6 using our 905 IDH^mut^ glioma culture.

### MSH6 KO induces TMZ resistance, which is overcome by KL-50

We next used our engineered MSH6-proficient and -deficient models to verify the previously reported role of MMR activity in mediating the antiproliferative and cytotoxic effects of TMZ in IDH^mut^ gliomas. As expected, lack of MSH6 increased TMZ resistance in all four models—MMR-:MMR+ GI50 ratios ranged from 2.2 to 30.4, with decreased TMZ cytotoxicity among all MSH6-deficient cells (**Fig. 2**, **Table 1**). In contrast, KL-50 not only retained activity against MSH6-deficient cells, but even suppressed proliferation, with BT142 and 905 also showing enhanced cell death (**Fig. 2**, **Table 1**). This suggests that KL-50 not only displays superior activity to TMZ in MMR-deficient glioma cells, but that MMR deficiency may further enhance KL-50 efficacy.

**Figure 2.**
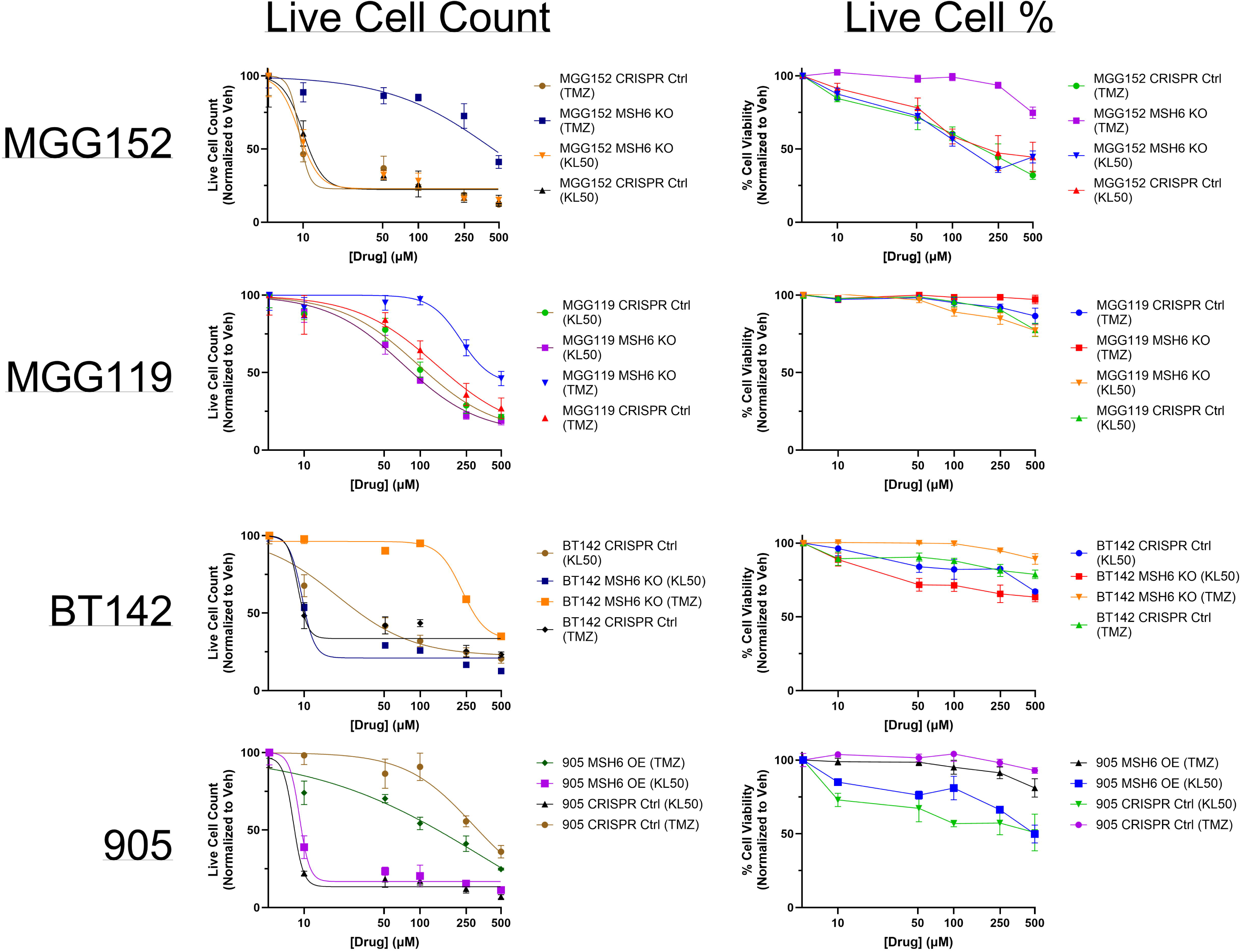
The effect of TMZ and KL-50 treatment on *in vitro* IDH^mut^ glioma cell growth and cytotoxicity in MMR-proficient and -deficient contexts using our MSH6-altered MGG152, MGG119, BT142, and 905 IDH^mut^ glioma cultures.

**Table 1.** Descriptive statistics associated with TMZ- and KL-50-mediated effects on cell growth and cytotoxicity in MMR-proficient and -deficient IDH^mut^ glioma cultures.

GI<sub>50</sub> and cytotoxicity by drug, cell line, and MMR status
| Drug | Cell line | GI<sub>50</sub> (μM) |  | GI<sub>50</sub> ratio<br>MMR–/MMR+ | Paired t-test, P |  |
| --- | --- | --- | --- | --- | --- | --- |
|  |  | MMR+ | MMR– |  | Live Cell Counts | Cytotoxicity |
| TMZ | MGG119 | 165.7 | 396.3 | 2.4 | **0.031** | 0.070 |
|  | MGG152 | 9.8 | 447.9 | 45.7 | **0.007** | **0.011** |
|  | BT142 | 9.9 | 300.9 | 30.4 | **0.015** | **0.005** |
|  | 905 | 142.3 | 317.4 | 2.2 | **0.020** | **0.019** |
| KL50 | MGG119 | 114.4 | 85.5 | 0.7 | **0.036** | 0.120 |
|  | MGG152 | 11.2 | 10.4 | 0.9 | 0.610 | 0.082 |
|  | BT142 | 30.0 | 10.3 | 0.3 | **0.010** | **0.009** |
|  | 905 | 9.5 | 8.3 | 0.9 | **0.035** | **0.032** |

Ratio = GI<sub>50</sub>(MMR–)/GI<sub>50</sub>(MMR+). Values and drug/cell-line order follow the source.
N=3 biological replicates per condition.
Bold P values indicate P < 0.05.

### KL-50 monotherapy induces apparent tumor eradication in a subset of mice engrafted with TMZ-naïve IDH^mut^ BT142 intracranial xenografts

To test whether KL-50 could work as a frontline therapy to treat IDH^mut^ glioma in a TMZ-naïve context, unmodified IDH^mut^ BT142 cells were orthotopically engrafted into NSG mice, followed by treatment with KL-50 or vehicle control. KL-50 extended median survival by 80% (vehicle=107.5 days vs. KL-50=194 days) (**Fig. 3A**). After 220 days post-engraftment the brains of the remaining long-term survivors were evaluated for residual tumor. As indicated by cellularity heatmaps, 2/8 KL-50 treated mice showed no evidence of residual tumor, with 1/8 showing equivocal tumor (**Fig. 3B**). In contrast, our previous KL-50 study involving MMR-proficient and -deficient IDH^wt^ GBM produced no long-term survivors or tumor eradication.(11) These data suggest that KL-50 is effective against IDH^mut^ gliomas in their MMR-proficient, TMZ-naïve state, and may be even more effective against IDH^mut^ gliomas than IDH^wt^ GBMs.

**Figure 3.**
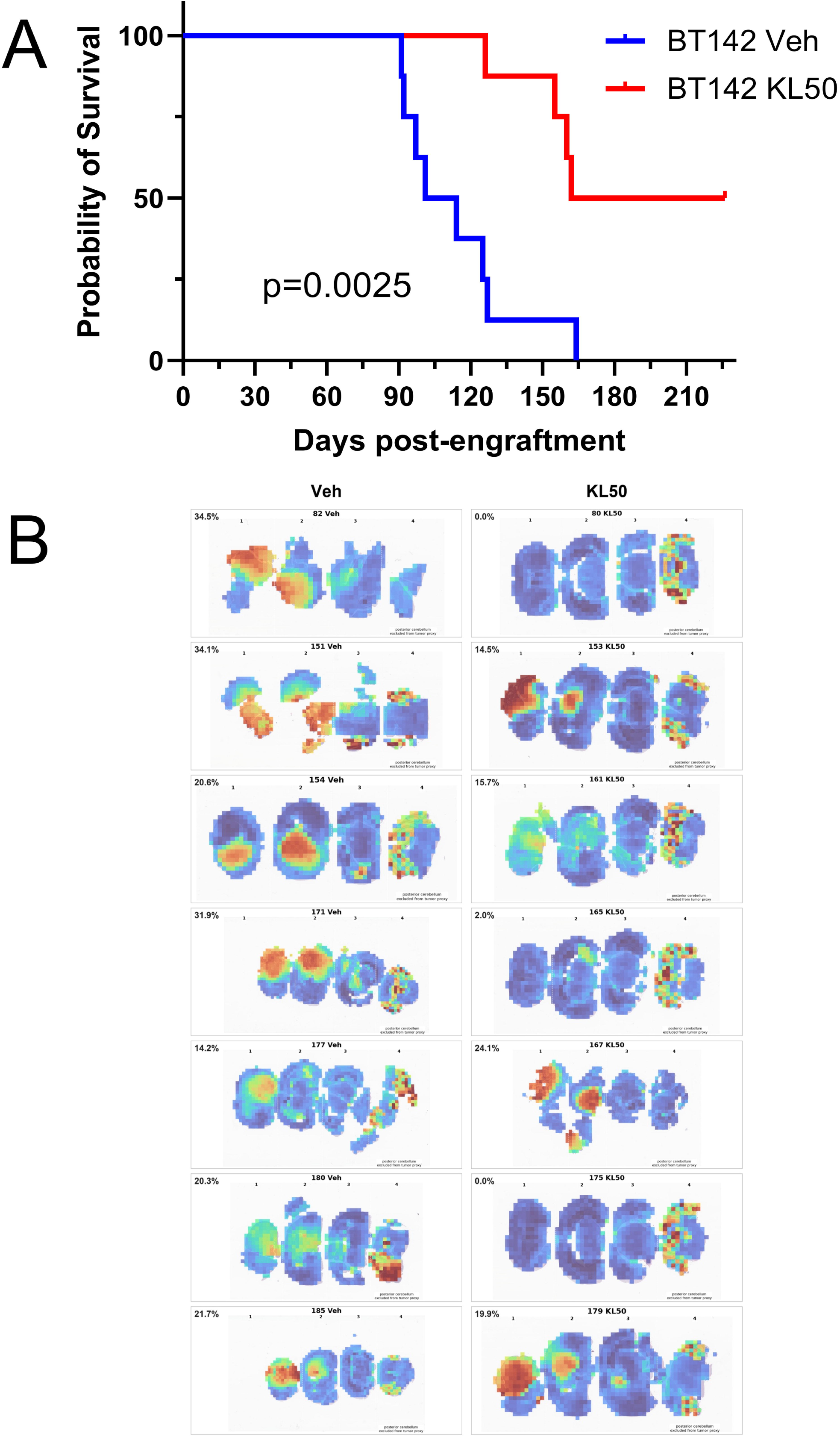
Investigation of KL-50 as a monotherapy using *in vivo* BT142 IDH^mut^ intracranial xenografts. (A) Kaplan-Meier survival analysis of TMZ-naïve BT142 intracranial xenografts treated with 10 mg/kg KL-50, M-F, for three weeks. (B) Histological H&E evaluation with a tumor cellularity heatmap describing residual tumor content in vehicle- and KL-50-treated mice.

### Combining KL-50 with the FDA-approved belinostat elicits a synergistic cytotoxic response

We and others previously showed that HDACi (panobinostat and belinostat) have unique epigenetic and transcriptomic effects on IDH^mut^ gliomas vs. IDH^wt^ GBM cells, suggesting that IDH^mut^ gliomas are particularly sensitive to HDACi.(13,24) Since monotherapies tend to be ineffective in glioma clinical trials, we sought to combine drugs that are particularly active against IDH^mut^ gliomas and potentially have synergistic mechanisms of action. Indeed, KL-50 plus belinostat showed synergistic toxicity against IDH^mut^ 905, BT142, and TS603 cells via LOEWE synergy analyses (**Fig. 4A-C**).

**Figure 4.**
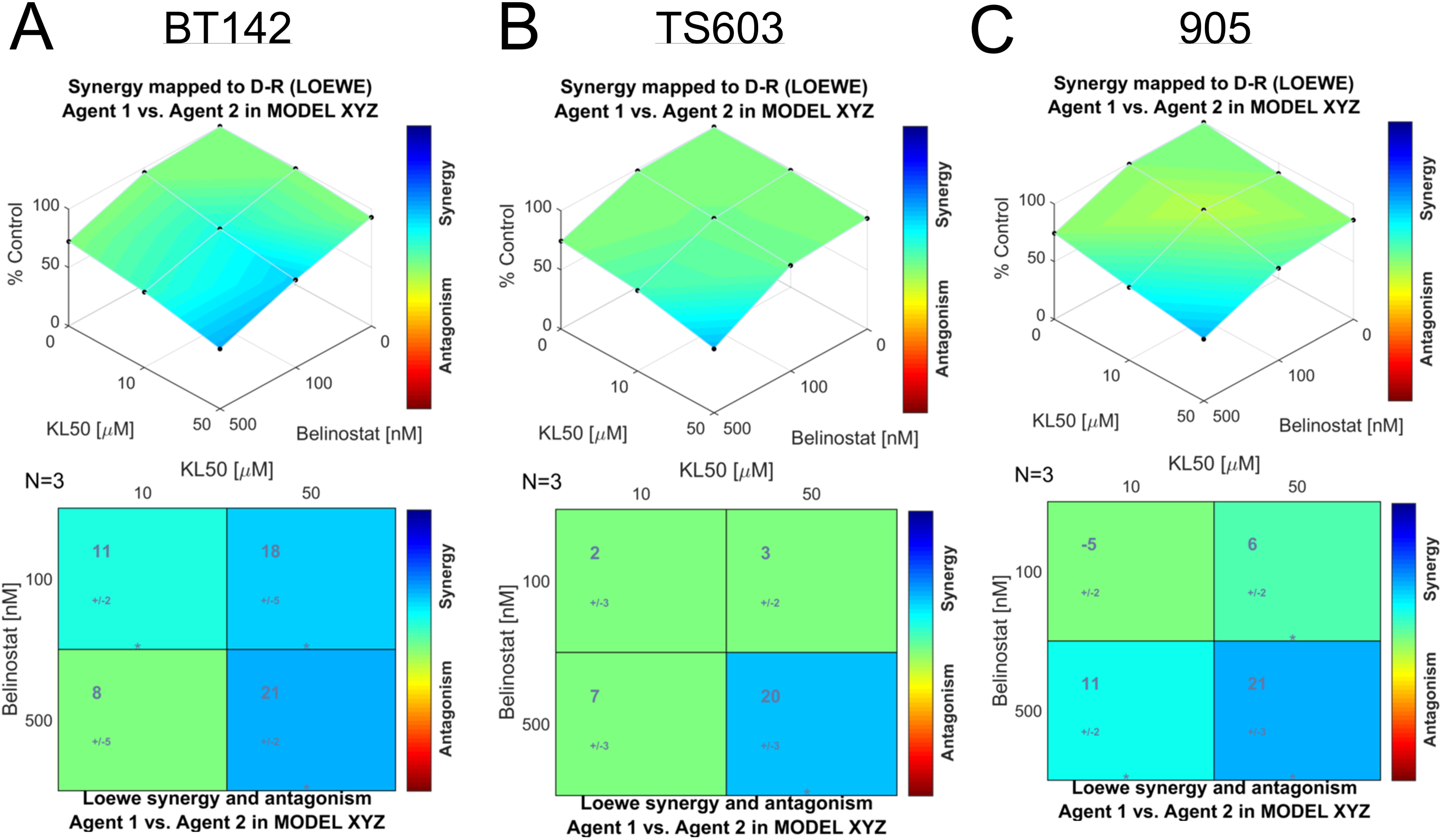
LOEWE synergy analysis of the combined cytotoxic effects of KL-50 and belinostat in BT142 (A), TS603 (B), and 905 (C).

Next, transcriptomic changes were analyzed to search for potential mechanisms of KL-50/HDACi synergy in IDH^mut^ glioma cells. IDH^mut^ glioma cultures were treated with panobinostat (10 nm) or belinostat (500 nm) for 24 hours, followed by RNA-Seq analysis. Gene ontology analysis showed that multiple DNA repair pathways were downregulated by panobinostat and belinostat, including Double Stranded Break Repair, Double Stranded Break Repair via Homologous Recombination, and Recombinational Repair, DNA Damage Response, DNA Repair, and Double Stranded Break Repair via Break-induced Replication (**Table 2**). This suggests ways in which HDACi could potentiate the ability of KL-50 ICLs to cause double-stranded breaks (DSBs) (**Table 2**). Highlighted downregulated genes involved in these gene ontology terms for both panobinostat and belinostat include *CHEK1*, *BRCA2*, *RAD51*, *FANCD2*, *FANCI*, and numerous *MCM* and *PARP* family genes.

**Table 2.** Gene ontology analysis of shared DNA repair and DNA damage response pathways downregulated in both our panobinostat- and belinostat-treated RNA-seq samples. panobinostat versus belinostat GO tables **7 shared GO terms; 91 exact shared symbols + 5 documented alias matches = 96 gene identities.** All displayed terms are enriched among downregulated genes in both datasets. Each FDR comes from its own source analysis; no new overlap P value was calculated. **Shared pathways and associated genes**

Shared pathways and associated genes
| GO term and ID | Exact genes | Shared genes* | Panobinostat FDR (BH) | Belinostat FDR (BH) |
| --- | --- | --- | --- | --- |
| double-strand break repair<br>GO:0006302 | 39 | 40 | $1.42 \times 10^{-11}$ | $1.36 \times 10^{-3}$ |
| Shared genes: *AP5S1; BRCA2; BRD9; BRIP1; CDCA5; DCLRE1A; DNA2; DTX3L; EPC1; FANCD2; FEN1; GINS2; HMGB2; HSF2BP; MCM2; MCM3; MCM4; MCM5; MCM6; MCM7; MCM9; MMS22L; MTA1; NSD2<sup>\*</sup>; NUCKS1; PARP2; PARP3; PARP9; PLK1; RAD51; RAD51AP1; RAD51D; RAD54L; RHNO1; RMI2; RNF138; SETMAR; SLF2; TOPBP1; TRIP13* |  |  |  |  |
| double-strand break repair via homologous recombination<br>GO:0000724 | 23 | 23 | $2.70 \times 10^{-7}$ | $3.36 \times 10^{-3}$ |
| Shared genes: *AP5S1; BRCA2; BRD9; EPC1; FEN1; GINS2; MCM2; MCM3; MCM4; MCM5; MCM6; MCM7; MCM9; MMS22L; NUCKS1; RAD51; RAD51AP1; RAD51D; RAD54L; RMI2; RNF138; SLF2; TOPBP1* |  |  |  |  |
| recombinational repair<br>GO:0000725 | 24 | 24 | $1.35 \times 10^{-8}$ | $4.13 \times 10^{-3}$ |
| Shared genes: *AP5S1; BRCA2; BRD9; EPC1; FEN1; GINS2; MCM2; MCM3; MCM4; MCM5; MCM6; MCM7; MCM9; MMS22L; NUCKS1; RAD51; RAD51AP1; RAD51D; RAD54L; RHNO1; RMI2; RNF138; SLF2; TOPBP1* |  |  |  |  |
| DNA damage response<br>GO:0006974 | 91 | 96 | $7.36 \times 10^{-16}$ | $5.02 \times 10^{-3}$ |
| Complete shared set (96 identities): *AP5S1; BARD1; BID; BRCA2; BRD9; BRIP1; CBX5; CDCA5; CHEK1; CIP2A<sup>\*</sup>; CTPS1; DCLRE1A; DDX11; DGCR8; DNA2; DONSON; DTL; DTX3L; E2F7; ENDOG; EPC1; EXO1; FAAP24; FAM111A; FANCD2; FANCI; FBXO5; FBXO6; FBXW4; FEN1; FOXM1; GINS2; GTSE1; HMGB2; HSF2BP; IFI16; KIF22; LIG1; MAP3K20<sup>\*</sup>; MAPK12; MC1R; MCM2; MCM3; MCM4; MCM5; MCM6; MCM7; MCM9; MLST8; MMS22L; MTA1; MUC1; NEIL1; NEIL3; NSD2<sup>\*</sup>; NUCKS1; PARP10; PARP2; PARP3; PARP9; PCLAF<sup>\*</sup>; PHLDA3; PIDD1; PLK1; POLD3; POLE; POLE2; PSMB5; RAD51; RAD51AP1; RAD51D; RAD54L; RBX1; RFC4; RHNO1; RMI2; RNASEH2B; RNF138; RRM1; RRM2; SET; SETD7; SETMAR; SLF2; SPINDOC<sup>\*</sup>; SUV39H1; TFAP4; TIMELESS; TOP2A; TOPBP1; TP53; TRIP13; UNG; WDR76; YAP1; ZBTB1* |  |  |  |  |
| DNA repair<br>GO:0006281 | 65 | 66 | $6.66 \times 10^{-15}$ | $5.08 \times 10^{-3}$ |
| Shared genes: *AP5S1; BARD1; BRCA2; BRD9; BRIP1; CDCA5; CHEK1; DCLRE1A; DNA2; DTX3L; EPC1; EXO1; FAAP24; FAM111A; FANCD2; FANCI; FBXO6; FEN1; GINS2; HMGB2; HSF2BP; KIF22; LIG1; MC1R; MCM2; MCM3; MCM4; MCM5; MCM6; MCM7; MCM9; MMS22L; MTA1; NEIL1; NEIL3; NSD2<sup>\*</sup>; NUCKS1; PARP2; PARP3; PARP9; PLK1; POLD3; POLE; POLE2; PSMB5; RAD51; RAD51AP1; RAD51D; RAD54L; RBX1; RFC4; RHNO1; RMI2; RNASEH2B; RNF138; RRM1; RRM2; SET; SETMAR; SLF2; TIMELESS; TOPBP1; TP53; TRIP13; UNG; ZBTB1* |  |  |  |  |
| double-strand break repair via break-induced replication<br>GO:0000727 | 7 | 7 | $5.39 \times 10^{-5}$ | $1.96 \times 10^{-2}$ |
| Shared genes: *GINS2; MCM2; MCM3; MCM4; MCM5; MCM6; MCM7* |  |  |  |  |
| signal transduction in response to DNA damage<br>GO:0042770 | 22 | 23 | $5.51 \times 10^{-5}$ | $3.59 \times 10^{-2}$ |
| Shared genes: *BID; BRIP1; CHEK1; DONSON; DTL; DTX3L; E2F7; FANCD2; FBXO6; FOXM1; GTSE1; MAP3K20<sup>\*</sup>; MAPK12; MUC1; PARP9; PIDD1; PLK1; RBX1; RHNO1; TFAP4; TIMELESS; TOPBP1; TP53* |  |  |  |  |

\* Shared counts include the five documented alias matches; Exact counts do not. An asterisk after a gene marks an alias match, not statistical significance (Table 3). The complete set of 96 shared identities appears under DNA damage response (GO:0006974). Genes recur across related terms, so pathway counts must not be summed.

Finally, a CRISPR KO viability screen with 905 IDH^mut^ glioma cells was performed to identify which gRNAs were depleted or enriched after KL-50 treatment—suggesting roles in KL-50 resistance or sensitivity, respectively. While none of the previously identified DNA repair genes were depleted or enriched after KL-50 exposure, the screen did identify KL-50 mediated depletion of gRNAs targeting *PKMYT1*, encoding the MYT1 kinase that blocks mitosis under stress from DNA damage (**Fig. 5A-B**).(25) *PKMYT1* is not only downregulated in IDH^mut^ glioma cells after HDACi (**Fig. 5C-D**), but its baseline expression is lower in TCGA IDH^mut^ astrocytomas and oligodendrogliomas vs. IDH^wt^ GBMs (**Fig. 5E**). These data provide additional mechanistic possibilities for the synergy between KL-50 and HDACi.

**Figure 5.**
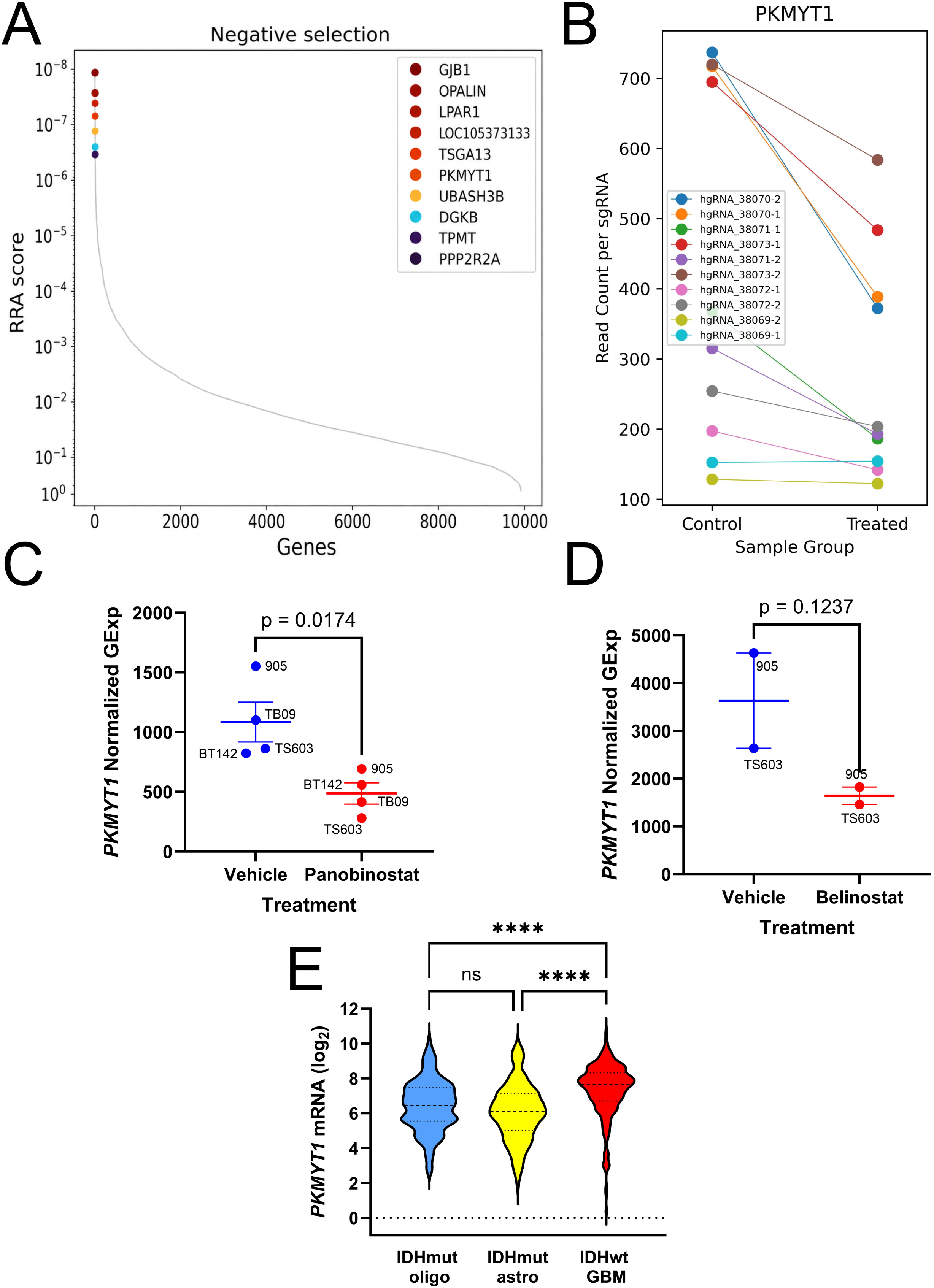
Identification of potential mechanisms of synergy between KL-50 and belinostat. (A) MAGECK analysis of our CRISPR KO screen data showing depleted genes in our 905 IDH^mut^ glioma culture after KL-50 treatment. (B) CRISPR KO screen data showing changes in *PKMYT1* gRNA after KL-50 treatment. (C) *PKMYT1* gene expression after panobinostat treatment of 905, BT142, TS603, and TB09 IDH^mut^ glioma cultures. (D) *PKMYT1* gene expression after belinostat treatment of 905 and TS603 IDH^mut^ glioma cultures (E) *PKMYT1* gene expression of sampled IDH^mut^ oligodendrogliomas, IDH^mut^ astrocytomas, and IDH^wt^ GBMs using the publicly available TCGA-GBMLGG dataset.

## Discussion

Acquisition of drug resistance remains a challenging problem in the effort to treat glioma patients. Therefore, it is critical to develop therapeutic strategies that can overcome known, and potentially unknown, resistance mechanisms. Here, using a panel of patient-derived IDH^mut^ glioma spheroid cultures, we show that KL-50, which was originally developed to overcome MMR-mediated TMZ resistance mechanisms in IDH^wt^ GBM,(10,11) is also effective against MMR-proficient and -deficient IDH^mut^ gliomas. MSH6 KO induces TMZ resistance in IDH^mut^ glioma cultures, just like what often happens in IDH^mut^ gliomas recurring after TMZ.(8,26) Owing to its ICL formation instead of 06-methylguanine-dependent anticancer activity, we confirmed that KL-50 does not depend on MMR pathways for its toxicity. In fact, KL-50 appears to be slightly *more* effective in MMR-deficient contexts. This suggests that MMR enzymes are not only necessary for TMZ-induced anticancer effects but also may play a role in repairing KL-50-mediated ICLs. Indeed, other reports suggest that MMR pathways facilitate ICL repair.(27–29) Further investigations into this are ongoing, but the data thus far indicate that KL-50 might be even more effective than expected against post-TMZ IDH^mut^ gliomas with acquired MMR deficiency.

Not only could ICL-forming agents like KL-50 overcome MMR-mediated TMZ resistance in recurrent tumors, but they might also be used as frontline adjuvant therapeutics. Such an approach would expand the target patient population and eliminate the generation of hypermutant, MMR-deficient recurrent IDH^mut^ gliomas. This hypermutant state accelerates tumor evolution and thereby acquisition of mutations associated with drug resistance during treatment.(8,26) Our data indicate that KL-50 is very effective against TMZ-naïve, MMR-proficient IDH^mut^ gliomas, even more so than in our previously published studies of IDH^wt^ GBM.(11)

Despite data showing its efficacy as monotherapy, other drug resistance mechanisms could eventually arise after KL-50. To circumvent this, we sought to explore its combination with a different class of chemotherapy, HDACi. Our previous work showed that IDH^mut^ gliomas are particularly sensitive to HDACi, and that HDACi causes more histone acetylation in IDH^mut^ gliomas than in IDH^wt^ GBM.(14,24) Considering the unique epigenomic background observed in IDH^mut^ gliomas, this suggests a unique and selective dependence of IDH^mut^ gliomas on HDAC epigenetic mechanisms.(13) Moreover, combining KL-50 and belinostat induced a synergistic cytotoxic response in IDH^mut^ gliomas.

Subsequent investigations into possible mechanisms of KL-50/HDACi synergy in IDH^mut^ gliomas revealed HDACi-mediated downregulation of DNA repair pathways, including genes involved in ICL or DSB repair such as *RAD51*, *CHEK1*, and *BRCA2*.(28,30) However, an unbiased CRISPR KO viability screen with KL-50 identified *PKMYT1*, a gene that was depleted in our screen and downregulated by HDACi. *PKMYT1* encodes MYT1 kinase, which initiates cell cycle arrest after DNA damage. Several reports have shown that loss of MYT1 sensitizes tumors to alkylating agents,(25,31,32) but MYT1 has not yet been connected to interstrand crosslink repair. Ongoing experiments are investigating this connection, as well as evaluating the novel KL-50/belinostat combination against *in vivo* models of IDH^mut^ glioma.

Together, this research advances the rationale for MGMT-sensitive ICLs, including KL- 50, as adjuvant therapy for IDH^mut^ glioma. This also suggests that KL-50 would be particularly useful in TMZ-treated recurrent due to potential role of MMR pathways in repairing KL-50 ICLs.(27–29) Finally, combining KL-50 with the FDA-approved HDACi belinostat may be a promising drug combination to potentiate tumor cytotoxicity, and could prevent the development of resistance mechanisms arising from either drug as monotherapy.

